# SenolyticSynergy: An Attention-Based Network for Discovering Novel Senolytic Combinations via Human Aging Genomics

**DOI:** 10.1101/2025.05.28.655258

**Authors:** Yaowen Ye, Dengming Ming

**Affiliations:** College of Biotechnology and Pharmaceutical Engineering, Nanjing Tech University, Nanjing 211816, China

**Keywords:** Senolytics, Aging Genome, Drug Combinations, Temsirolimus, Nitazoxanide, Anti-Aging Interventions

## Abstract

Senolytics, a category of drugs targeting aging processes, have garnered significant attention since their emergence in 2015. Unlike traditional drug development approaches that rely on randomized screening, research on aging-related pharmaceuticals has employed mechanism-based strategies, resulting in the discovery of the pioneering combination therapy of dasatinib (D) and quercetin (Q). Although preliminary studies with senolytic drug combinations have shown promising outcomes, the predictive capabilities of the research in this field remain limited by the extensive experimental data requirements. In this study, we employed differential gene expression analysis and machine learning techniques to investigate the combinatorial effects of senolytic drugs. We identified 1624 core aging-related genes and used this dataset to retrain a multimodal attention mechanism model, creating a specialized framework: SenolyticSynergy for predicting effective senolytic drug combinations. We then utilized 63 established senolytic compounds as starting points for combination testing, developing a comprehensive dataset of 1953 potential drug combinations for aging interventions. Following rigorous filtration, we identified 190 high-confidence drug combinations and predicted their synergistic scores. Among these combinations, ten demonstrated exceptionally high synergistic scores, exceeding 8. The combination of temsirolimus and nitazoxanide ranked first and may be the most promising candidate. The analysis of literature data and computational studies of molecular structures using 3D modeling validated the accuracy of these predictions. This framework paves the way for large-scale research into anti-aging drug combinations, advancing research capabilities in this field.

**GRAPH ABSTRACT:** 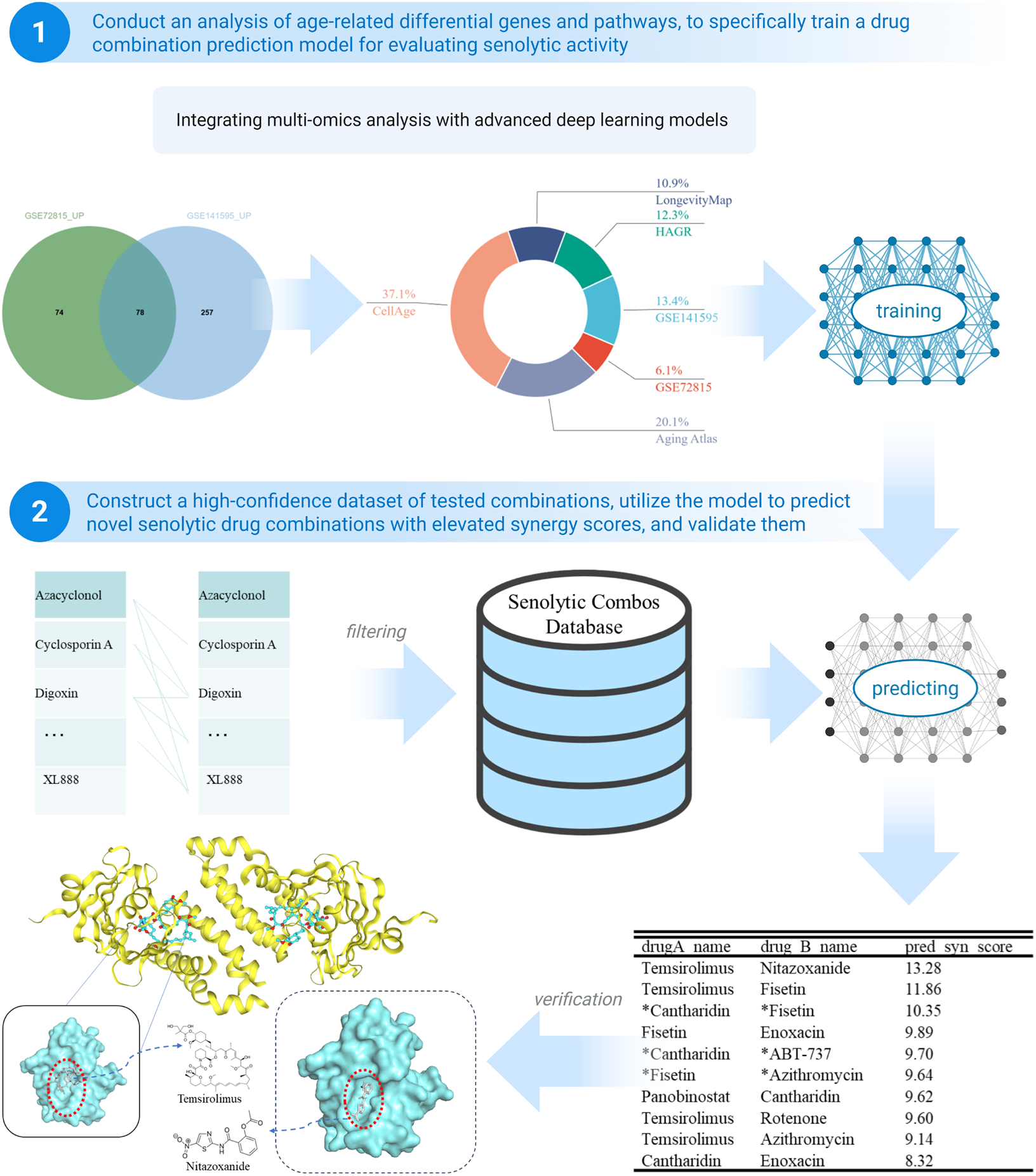

## INTRODUCTION

In recent years, the scientific community has redefined aging as a chronic disease involving complex biological processes (López-Otín et al., 2023). Its pathological trajectory is intertwined with the mechanisms of malignant tumors(Hanahan, 2022) and the pathological development of neurodegenerative diseases. Within this macro- physiological process, the biological homeostasis of cells and tissues is affected, leading to a gradual decline in cellular function and genomic stability. Together, these two factors increase the risk of cancer and neurodegenerative diseases as we age. If aging is viewed as a disease, senescent cells are the primary targets for intervention. Senolytics are a diverse class of drugs specifically targeting aging cells for clearance. Using combinations of senolytic drugs, it is possible to delay and inhibit aging-related behaviors at various levels of cells, tissues, or organs, ultimately intervening in the aging process of individuals and achieving healthy aging.

In the past decade, numerous machine learning-based models have been developed to predict drug combination synergy. For instance, Zhao and colleagues proposed the Dual Feature Fusion Network for Drug-Drug Synergy prediction (DFFNDDS), which utilizes a fine-tuned pre-trained language model and dual feature fusion mechanism to predict synergistic drug combinations(M. Xu et al., 2023). Wang and colleagues introduced EDDINet4, which enhances drug-drug(H. Wang et al., 2025) interaction (DDI) prediction via Information Flow and Consensus-Constrained Multi-Graph Contrastive Learning for precise DDI prediction. Li and colleagues built a two-view deep learning model, JointSyn(X. Li et al., 2024), for predicting the synergistic effects of drug combinations and applied it to find drug combinations for pan-cancer. Monem and colleagues proposed MultiComb (Monem et al., 2024), a multi-task deep learning (MTDL) model designed to predict the synergy and sensitivity of drug combinations simultaneously. This model creatively utilizes a graph convolution network to represent two drugs’ Simplified Molecular-Input Line-Entry (SMILES), generating their respective features. Guo and colleagues introduced SynergyX, a multi-modality mutual attention network to improve anti-tumor drug synergy prediction (Guo et al., 2024), which dynamically captured cross-modal interactions, allowing for the modeling of complex biological networks and drug interactions.

This study aimed to analyze the differential gene expression profiles between young and elderly populations and identify gene fragments closely related to aging. These gene fragments serve as key input elements for the drug-gene attention mechanism model, which predicts and optimizes the potential synergistic benefits of different anti-aging drug combinations. This study provides scientific evidence for precision medicine.

In this study, we utilized various R packages and methods(Tang et al., 2023) to analyze and process GSE141595(Hickson et al., 2019) and GSE72815(Farr et al., 2015) to accurately identify significantly differentially expressed gene fragments between the two groups. Subsequently, we adopted the attention-based mechanism network. We integrated the identified aging-related differential genes as a targeted subset for model training, focusing on developing a drug combination prediction model for aging-related diseases to achieve a precise prediction of synergistic effects between drugs.

We successfully identified multiple core differential genes related to aging through differential analysis, including PKP1, NRAP, and CMA1. For model fitting pre-screening, we employed an approach that involved the initial screening and subsequent calculation of 1953 drug combinations formed by 63 senolytics discovered and experimentally validated by July 1, 2024. After the screening, 193 valid entries were retained for drug combination synergy scoring. Ultimately, we retrained the prediction model to obtain ten drug combinations, including tsirolimus + nitazocine and tsirolimus + fiserone, which displayed exceptionally high synergistic scores in the model evaluation (pred_syn_score > 8), indicating significant therapeutic effects in aging interventions. Drugs with synergy scores exceeding eight were selected for literature validation.

Clustering analysis of differential genes has emphasized the significance of aging-related core pathways, such as hsa04020, demonstrating a robust theoretical foundation for a deeper understanding of the pathological mechanisms of aging diseases. Furthermore, published experimental studies have confirmed the synergistic effects of several drug combinations predicted by this model. For example, a survey by Ren(Ren et al., 2016) indicated that the combination of Cantharidin + ABT-737, among others, significantly improved the clearance of aging cells while maintaining low toxicity to normal cells. This discovery expands the horizons of anti-aging treatment strategies. This validates the feasibility of machine learning in discovering drug combinations, offering a more promising therapeutic outlook than single-drug treatments.

## MATERIALS AND METHODS

### Youth-Old Age Differential Gene Expression Dataset

Microarray data for this study were retrieved from the National Center for Biotechnology Information (NCBI) with accession numbers GSE72815 and GSE1415955. In the GSE72815 study, 58 healthy women were included, with 19 in the young women group (mean age ± standard deviation: 30.3 ± 5.4 years), 19 in the elderly women group (73.1 ± 6.6 years), and 20 older women (70.5 ± 5.2 years) who received 3 weeks of estrogen (E) treatment. Based on widely accepted criteria (false discovery rate [q] < 0.10), aging influenced a total of 678 genes and 12 pathways, including a subset of genes known to regulate bone metabolism.(Farr et al., 2015) GSE141595 involves gene profiling of trabecular bone biopsies in postmenopausal women who received either placebo or denosumab treatment for 3 months and gene analysis in young women who did not undergo treatment. We specifically utilized data from two groups: postmenopausal women receiving a placebo and young women not undergoing treatment.(Hickson et al., 2019)

### Senolytics Dataset

We obtained part of our single-drug dataset from Smer-Barreto(Smer-Barreto et al., 2023), then added the newest and found senolytics as supplementary to make the final table as follows:

**Table 1.**
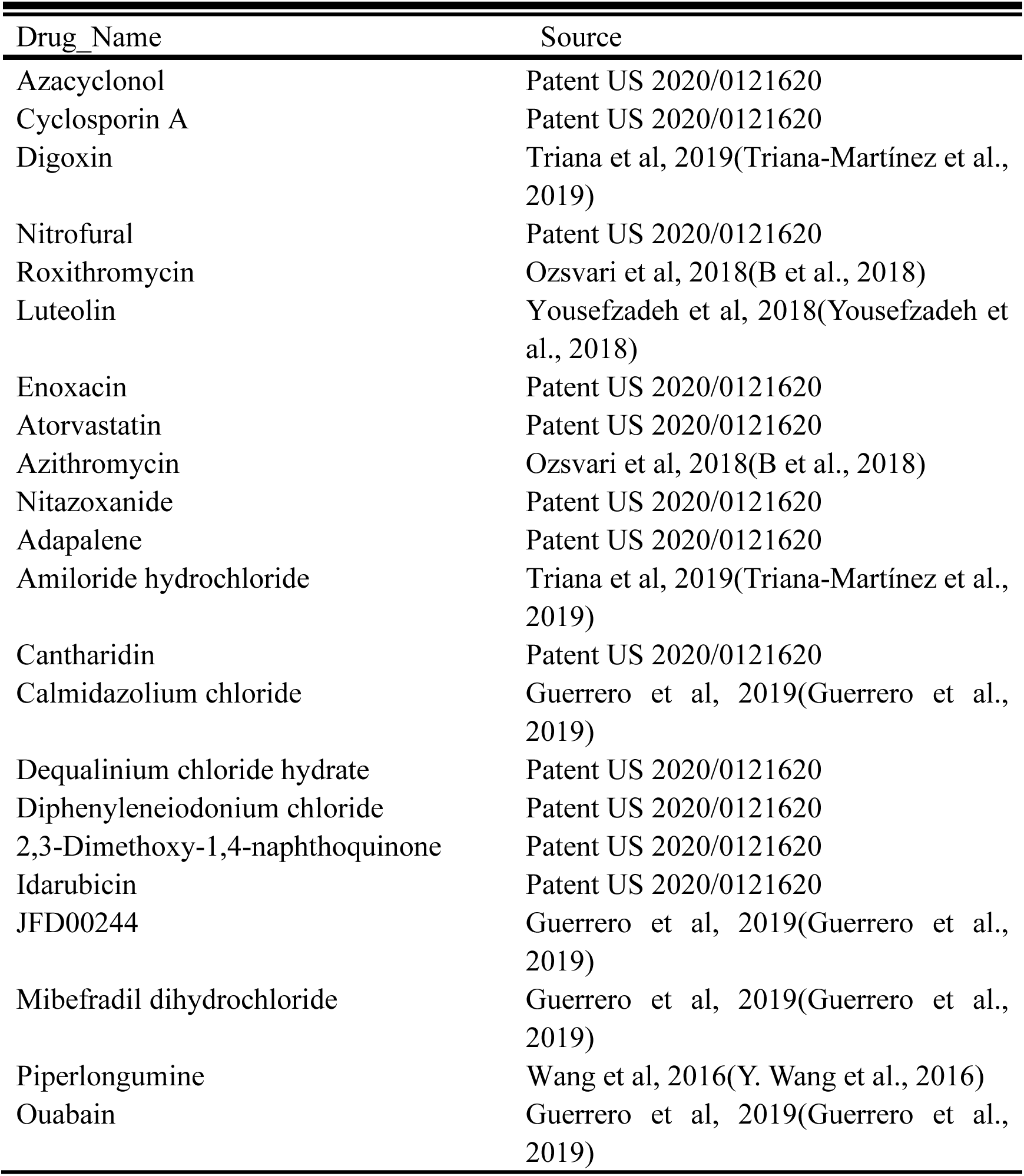
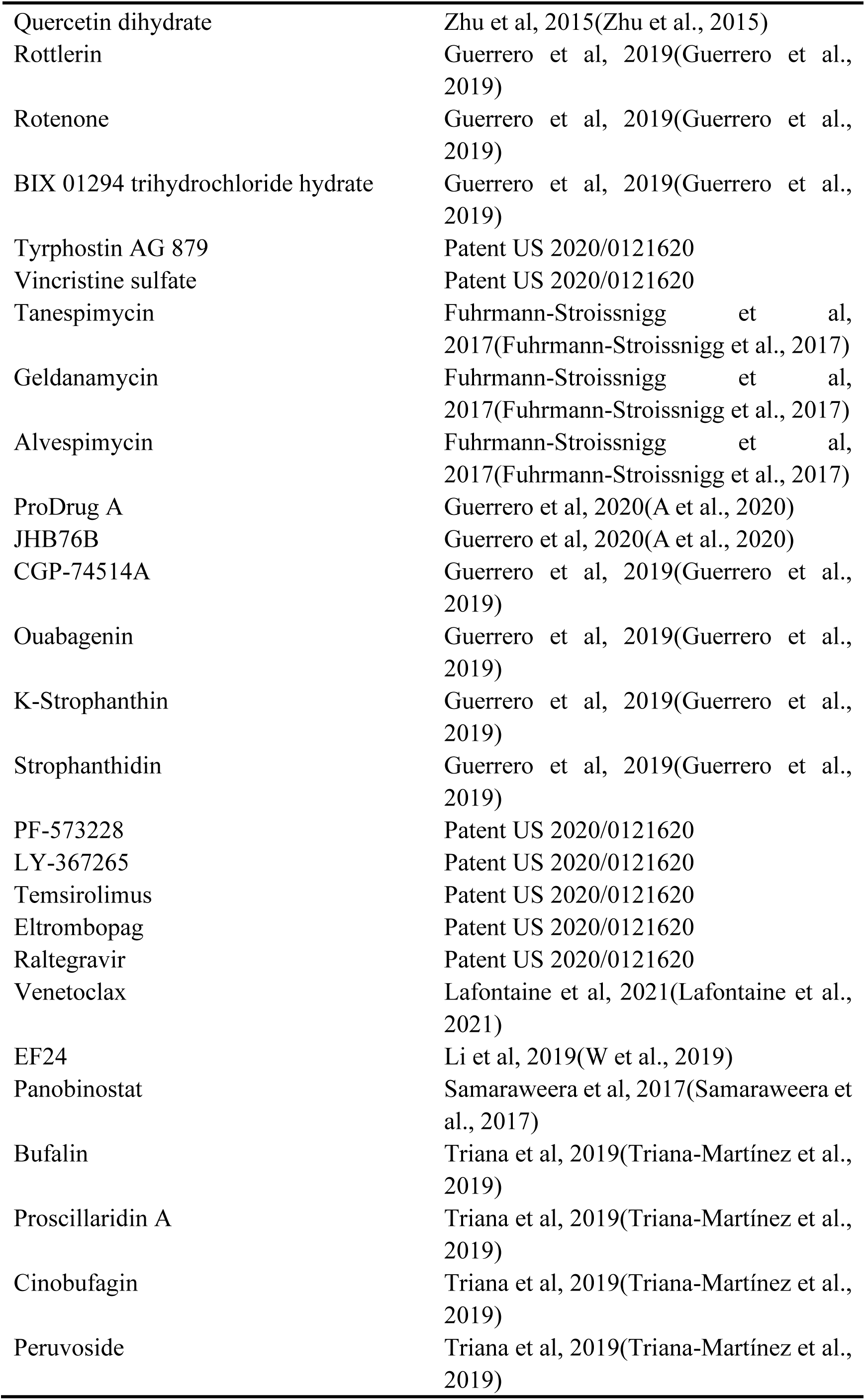
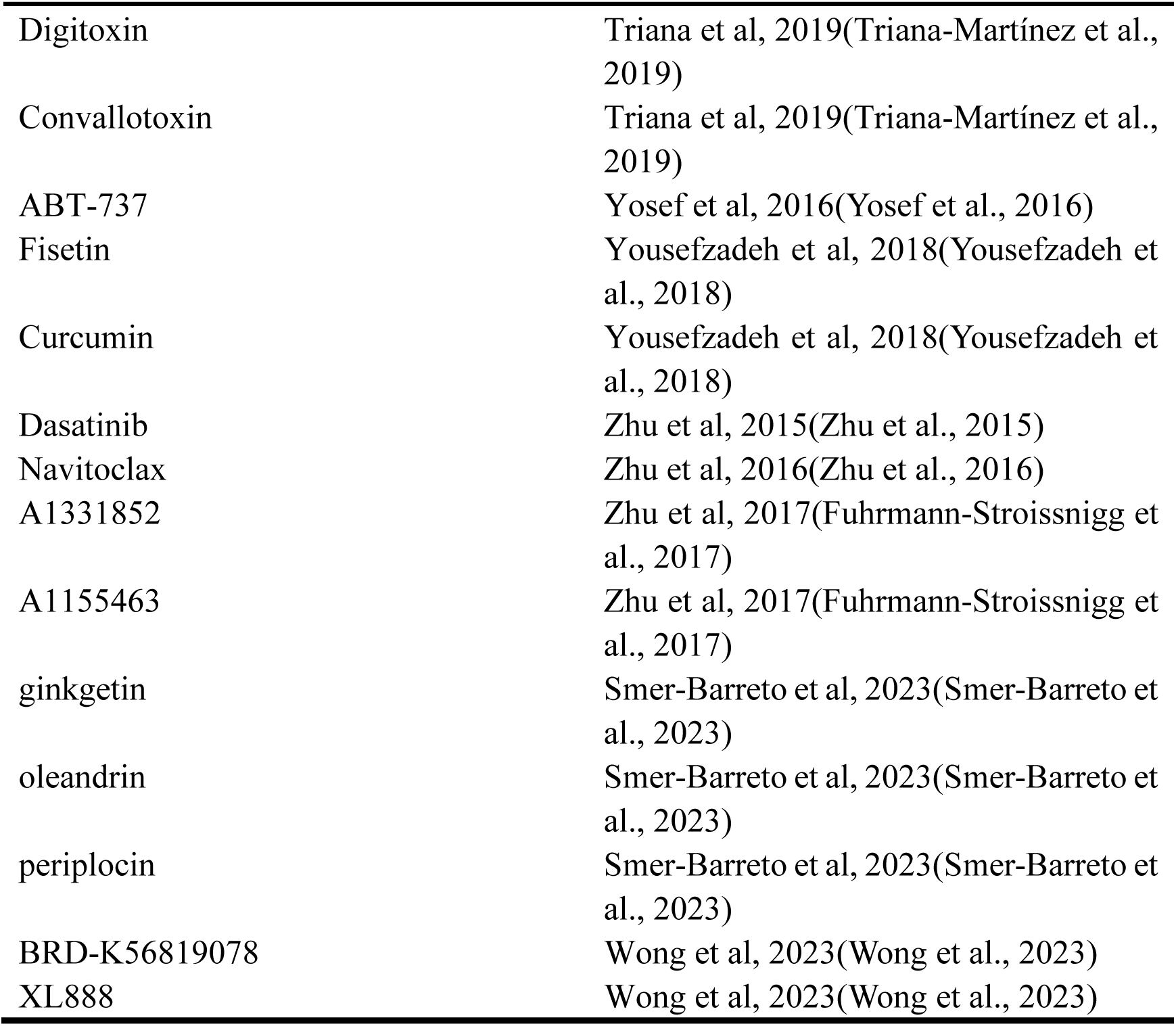
Single senolytic summary sheet (till July 1st, 2024).

### Aging-Related Target Gene Dataset

The target genes were selected from the following six sources:

1. From the differential gene analysis of GSE72815, 152 entries with logFC>1 and 11 entries with logFC<- -1 were selected.
2. From the differential gene analysis of GSE141595, 335 entries with logFC>1 and 90 with logFC<-1 were selected.
3. The Human Ageing Genomic Resources (HAGR) database(de Magalhães et al., 2024), specifically the GenAge resource package (https://www.genomics.senescence.info/genes/index.html), was used to download the latest stable version of human aging-related genes (https://www.genomics.senescence.info/genes/human_genes.zip). A total of 307 gene entries were extracted from GenAge and designated as Source Three in the tables, named GenAge_human.
4. The latest stable version of the LongevityMap(Budovsky et al., 2013)(https://www.genomics.senescence.info/longevity/) was obtained from the LongevityMap(https://www.genomics.senescence.info/longevity/longevity_genes.zip), reflecting the current understanding of human longevity genetics. However, this database includes records of negative results; therefore, only 273 gene entries with "significant" status in the Association column were selected and recorded in the longevityMap table.
5. A list of genes associated with cellular senescence was obtained from the CellAge database26 (https://genomics.senescence.info/cells/), which focuses on cellular senescence genes (https://genomics.senescence.info/cells/cellAge. zip). Entries under the "Unclear" attribute of the Senescence Effect were excluded, and the remaining 927 records were extracted and recorded in the table.
6. From the Aging Atlas database(“Aging Atlas,” 2021), aging-related gene sets were selected for download within the Aging-related gene sets section, resulting in 503 entries recorded in the Aging Atlas table.

The total number of gene entries under these six approaches was 2,566. After removing duplicate entries, 1,769 entries remained for analysis. Finally, gene ID conversion was performed, and entries that could not be converted were removed, resulting in the final selection of 1,624 target genes.

**Figure 1.**
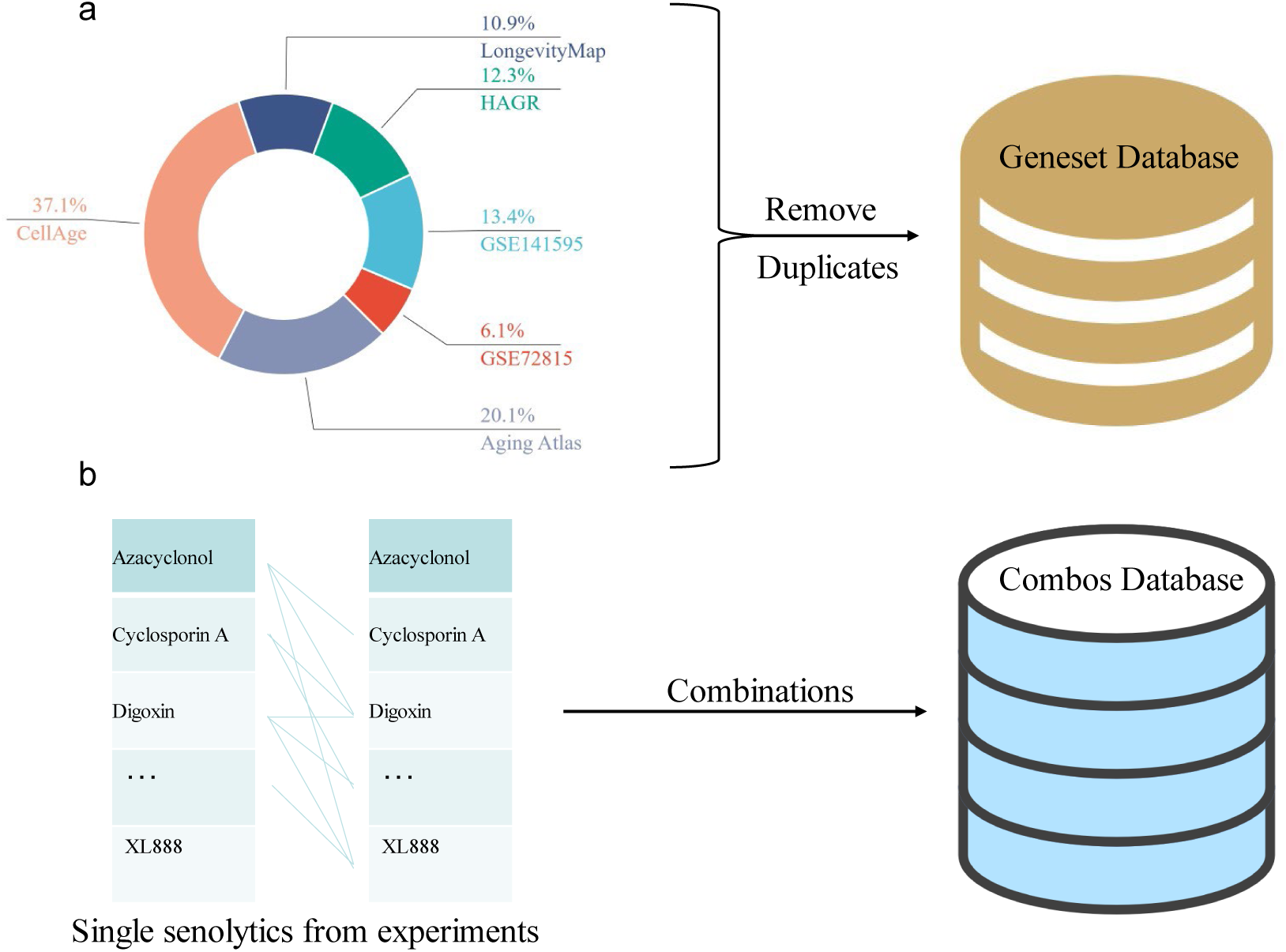
Database Components. (a) This pie chart depicts the distribution of different databases in the entire dataset. The most significant proportion is occupied by "CellAge" at 37.4%, followed by "Aging Atlas" at 20.1%, "GSE147591" at 13.4%, "Longevity Map" at 10.9%, "HAGR" at 12.3%, and "GSE72815" at 6.1%. After preprocessing, a 1624 unique gene entries database was obtained by removing duplicates from the remaining data. (b) The left column presents the drugs used in individual experiments, such as Cyclosporin A and Digoxin. The drugs were combined pairwise to create a new database, resulting in 1953 unique drug combinations.

### Senolytics Combination Efficacy Prediction Model

Regarding model prediction, we introduced SenolyticSynergy, a classical attention-based(Vaswani et al., 2017) mechanism network designed to enhance the prediction of the synergistic effects of senolytic drugs. This model utilizes a classic attention structure that effectively integrates multiple omics data within its framework.

**Figure 2.**
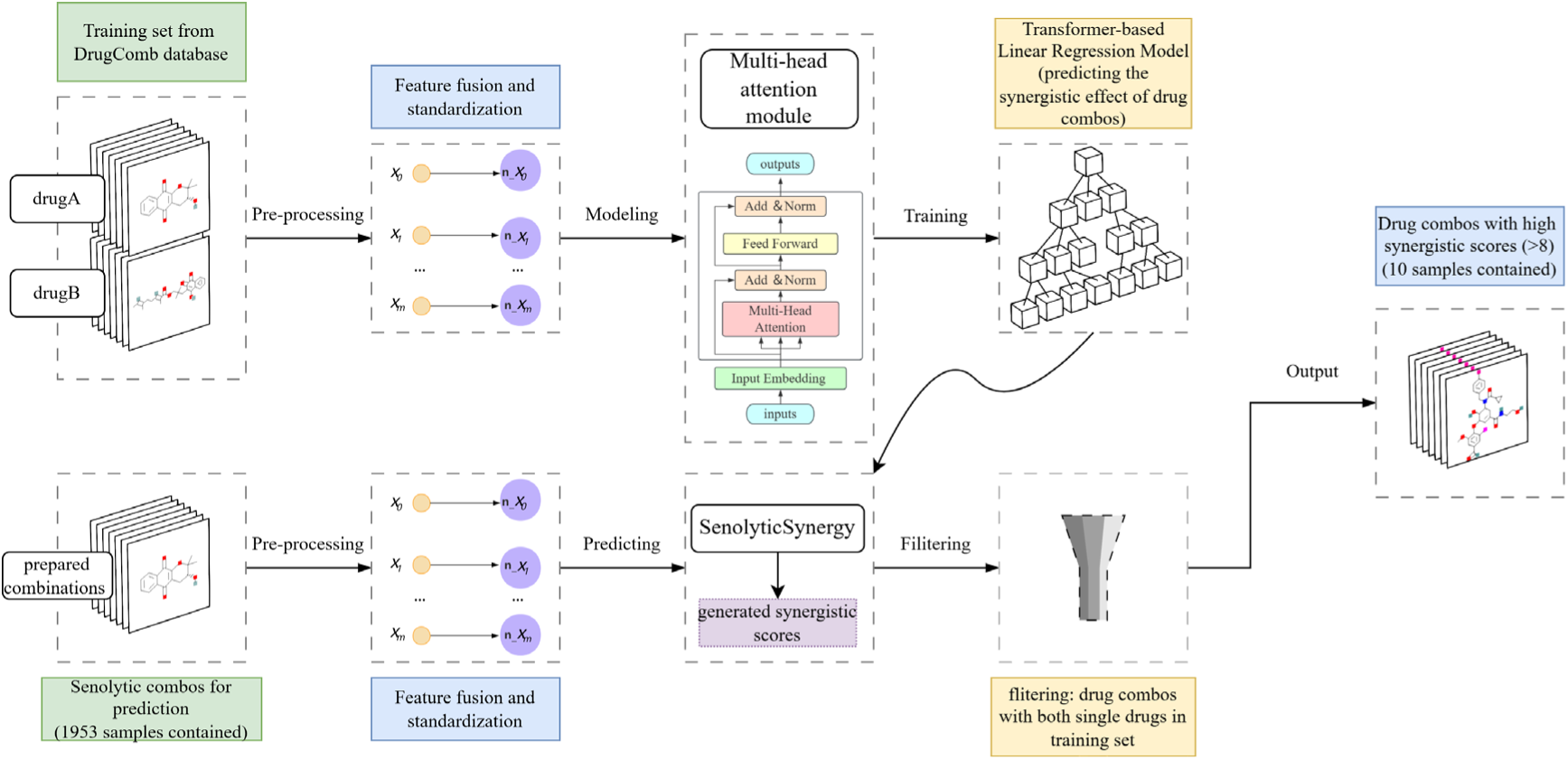
The overview of SenolyticSynergy.

The training set collects 330,917 drug combinations from the DrugComb database consisting of 354 single drugs and 170 cell lines with S synergy scores(Malyutina et al., 2019), and performs drug-target gene feature fusion and feature standardization during preprocessing, builds the model with the multi-attention mechanism and inputs the features for training, and selects the optimal parameters as our prediction model: SenolyticSynergy. 1953 senolytic combinations used for prediction were also subjected to feature fusion and standardization, and the corresponding synergy scores of each combination were predicted by the SenolyticSynergy model. Meanwhile, we further filtered out 190 cases of senolytic combinations in which all single drugs appeared in the training set, and selected those combinations with synergy scores greater than 8 as the final output, based on the theory of Lin et al. that the training set contained single drugs within the prediction set that improved the accuracy of synergy prediction of drug combinations (40% to 90%)(Lin et al., 2022).

## RESULTS

### Differential Genetic Analysis

We obtained the gene expression data files of the GSE141595 and GSE72815 datasets by accessing the GEO database and conducting differential gene expression analysis. We applied |log2FC| greater than 1.0 and Padjust less than 0.05 as filtering criteria, and the outcomes are shown in Figure 3. Furthermore, we identified the core genes by performing an intersection analysis between GSE141595 and GSE72815 (Table 2).

**Figure 3.**
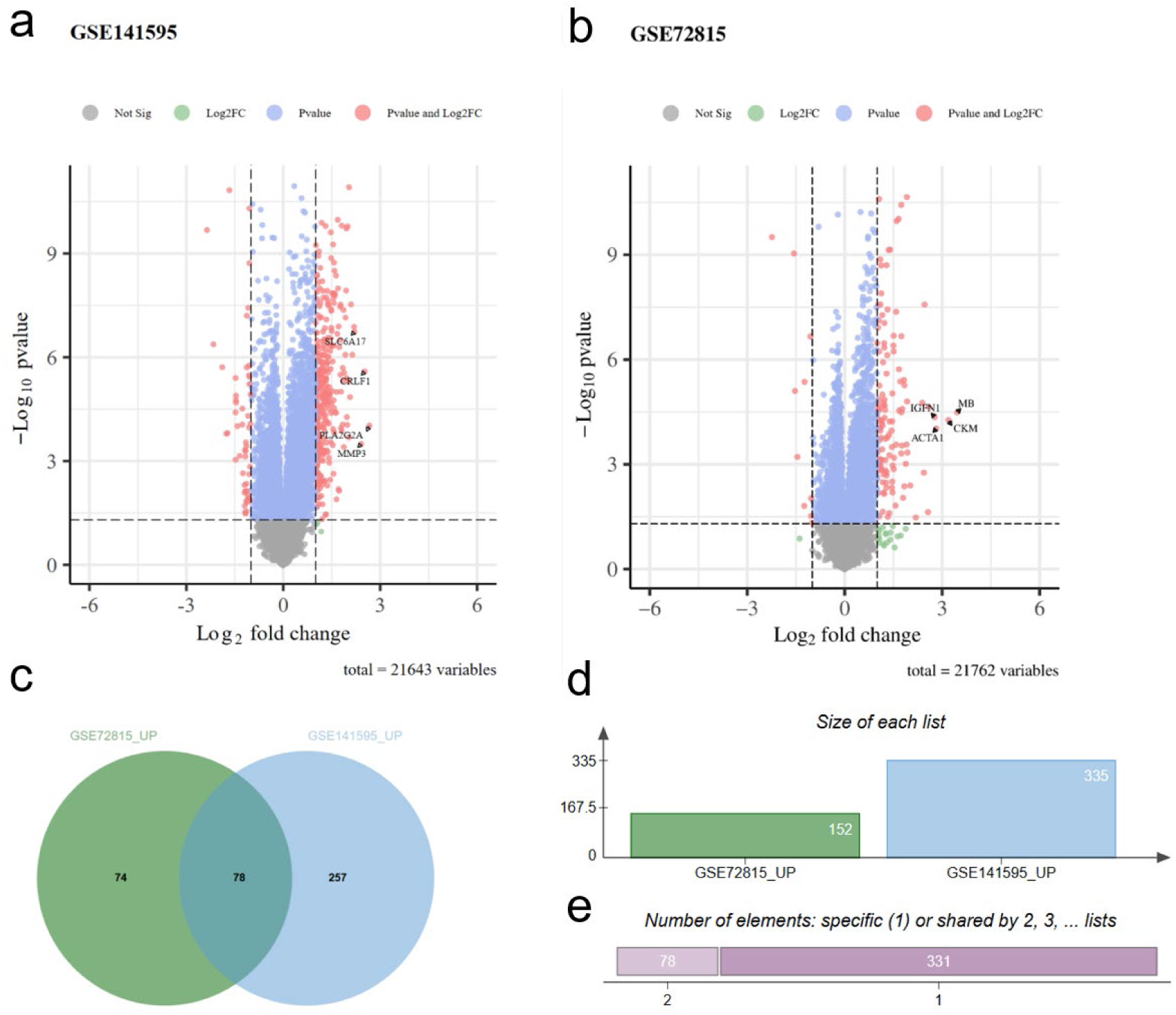
Aging Genome Analysis plots (a, b) illustrating the differentially expressed genes in the GSE141595 and GSE72815 datasets. The horizontal axis represents the log2 fold change, and the vertical axis represents the -log10 p-value. Each point represents a gene, with color coding and labels denoting the gene’s significance based on thresholds of fold change and p-value. In GSE141595, the significantly upregulated genes included SLC6A17, CRLF1, PLAG2A, and MMP3. Similarly, in GSE72815, the significantly upregulated genes included MB, CKM, IGFN1, and ACTA1. (c) displays the overlap of upregulated genes between the two datasets, revealing 78 genes commonly upregulated in the intersection of GSE141595 and GSE72815. This significant overlap mutually validates the reliability of the differential analysis data for the two gene sets. (d, e) present the total number of upregulated genes in the two datasets, amounting to 409 after excluding duplicate entries.

**Table 2.**
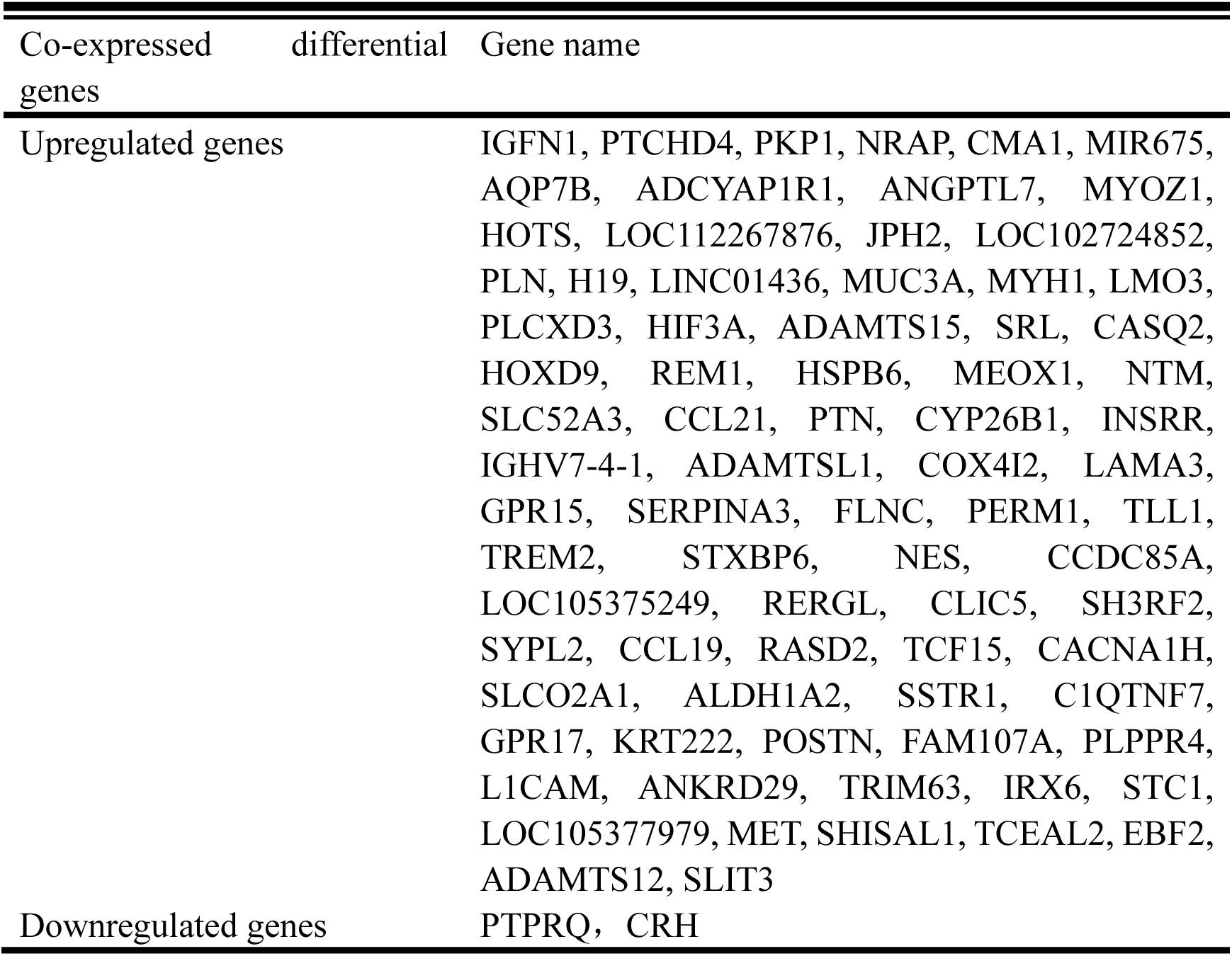
common DEGs from GSE141595 and GSE72815.

The genes listed in Table 2 have demonstrated high relevance to aging. For illustrative purposes, we selected five genes for discussion. H19(Xia et al., 2024) is a long non-coding RNA (lncRNA) that regulates gene expression and cellular aging. HIF3A encodes hypoxia-inducible factor 3α, a key regulatory factor in the cellular response to low-oxygen environments relevant to aging and lifespan regulation(Jaskiewicz et al., 2022). The TREM2 gene is associated with neurodegenerative diseases, such as Alzheimer’s disease, and influences the aging process of the brain(Jay et al., 2017). SERPINA3, which encodes α1-antichymotrypsin, is associated with inflammation and aging(Sánchez-Navarro et al., 2021). Lastly, ADAMTS12 encodes a protease involved in extracellular matrix remodeling associated with tissue aging and repair(El Hour et al., 2010).

### Enrichment analysis

Enrichment analysis is a critical method in bioinformatics, primarily employed to elucidate the functional tendencies of gene sets within specific biological contexts(Subramanian et al., 2005). GO enrichment analysis utilizes the GO database for functional annotation and enrichment analysis of given gene sets. The GO database categorizes genes into three primary annotation categories: Biological Process (BP), Cellular Component (CC), and Molecular Function (MF).

### Pathway analysis

The KEGG (Kyoto Encyclopedia of Genes and Genomes) pathway enrichment analysis is a significant bioinformatics method(Kanehisa & Goto, 2000) for investigating genes’ involvement in specific biological processes and their participation in complex pathways, including cellular metabolism and signal transduction. This study utilized the R packages clusterProfiler and pathview to perform KEGG pathway enrichment analysis of the extracted core aging-related genes.

The table illustrates pathways involving genes that occur two or more times. There are 11 significant signaling pathways, including the calcium signaling pathway, which concerns the transmission of signals by calcium ions within cells. Calcium ions play crucial roles in organisms, including cell growth, differentiation, apoptosis, and muscle contraction.

Most of the signaling pathways presented in this table are intricately linked to aging. For instance, hsa04020, which ranks highest in relevance, involves calcium ions as vital cellular messengers in various biological functions, including cell proliferation, differentiation, programmed cell death, and muscular contractions. This pathway comprises two primary components: (1) External calcium ion sources: Cells obtain calcium ions from their surroundings through diverse calcium channels located on the cellular membrane, such as voltage-gated channels (VOCs), receptor-gated channels (ROCs), and store-operated channels (SOCs). (2) Internal calcium ion sources: The endoplasmic reticulum/sarcoplasmic reticulum (ER/SR) functions as a critical calcium ion storage site, where inositol 1,4,5-trisphosphate receptors (IP3Rs) and ryanodine receptors (RYRs) control the release of calcium ions.

Furthermore, the core pathways in Table 3 include proteins and signaling factors highly relevant to cancer regulation, such as the p53 and Wnt signaling pathways within the MAPK signaling pathway (hsa04010). The Wnt signaling pathway is critical for embryonic development, tissue repair, and cancer, influencing cell behavior by regulating proliferation, differentiation, migration, and stemness maintenance(Clevers, 2006). Abnormal activation of the Wnt pathway is associated with various cancer types, including colorectal and liver cancers, with mutations in tumor suppressor genes in the Wnt pathway (such as APC and AXIN2) leading to overactivation of β-catenin and promoting the growth and self-renewal of tumor cells.

**Table 3.**
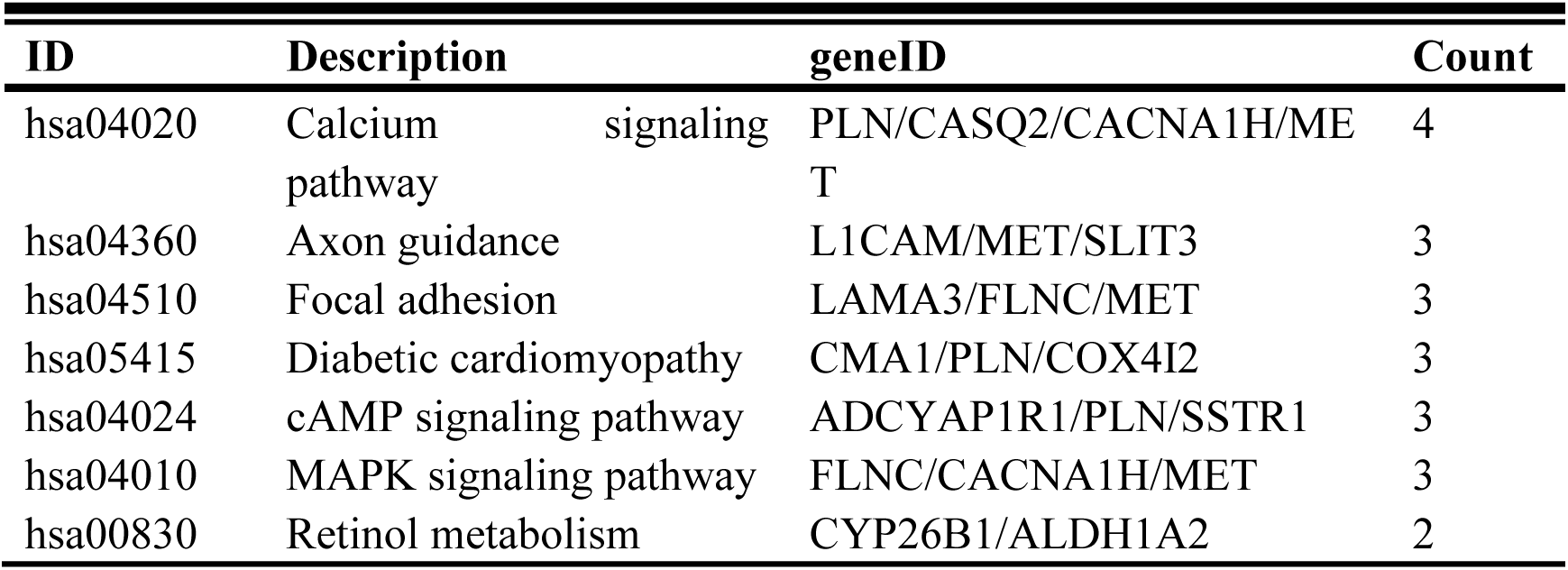
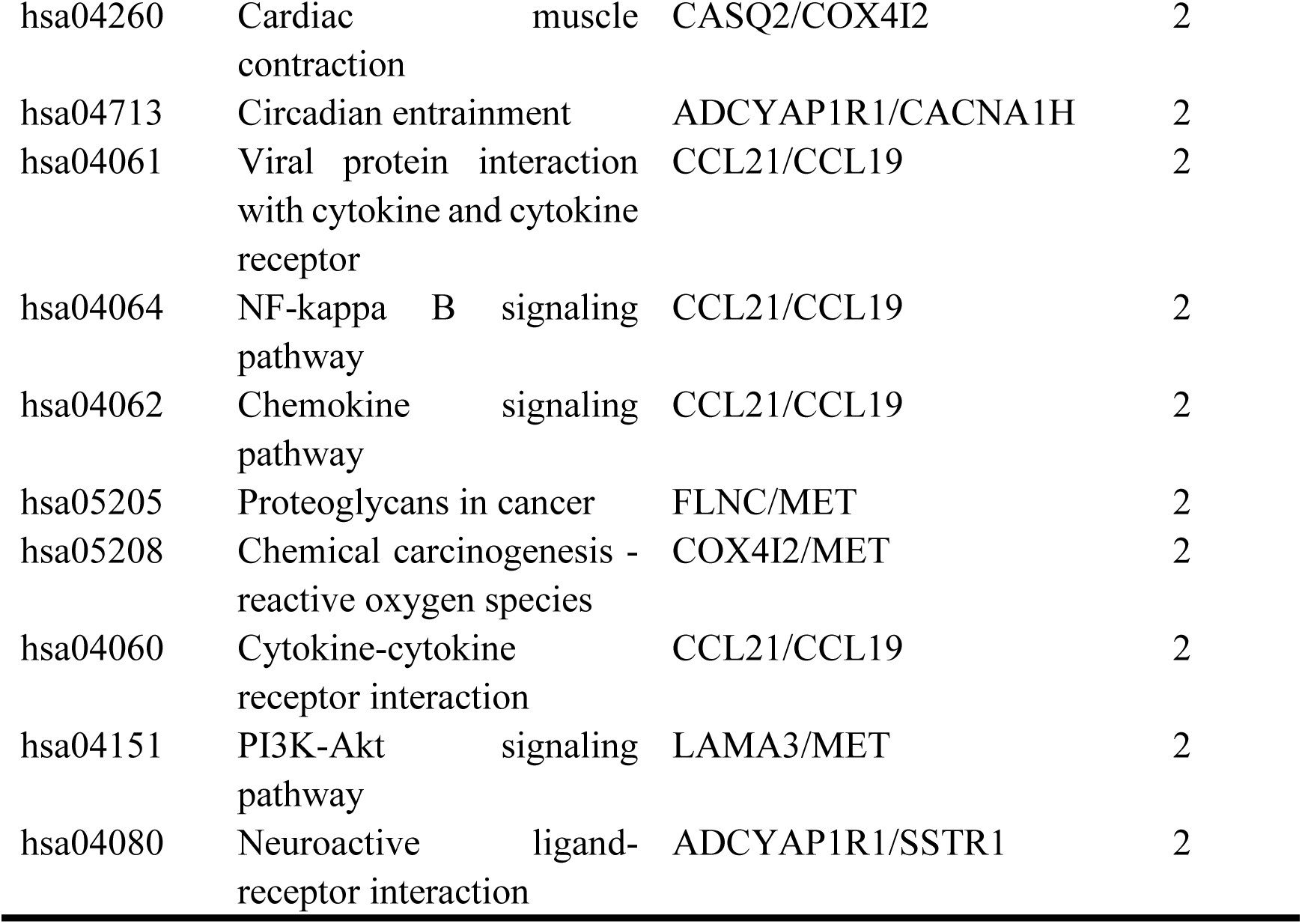
Enrichment pathway list of aging genome DEGs.

### Synergy Prediction and Interpretation

The prediction model we employed is called SenolyticSynergy. In this study, we integrated the dataset of omic genes 1624 genes to features and made the model specifically for senolytic target utilization. The following Table 4 presents the prediction results sorted by the prediction synergy score.

**Table 4.** Drug combinations with the top synergistic scores (syn_score > 8)

| drugA_name | drug_B_name | pred_syn_score | available reference |
| --- | --- | --- | --- |
| Temsirolimus | Nitazoxanide | 13.28 | / |
| Temsirolimus | Fisetin | 11.86 | / |
| *Cantharidin | *Fisetin | 10.35 | Frezzato et al.<br>(Frezzato et al.,<br>2020) |
| Fisetin | Enoxacin | 9.89 | / |
| *Cantharidin | *ABT-737 | 9.70 | Ren et al. (Ren et<br>al., 2016) |
| *Fisetin | *Azithromycin | 9.64 | Shao et al. (Shao<br>et al., 2024) |
| Panobinostat | Cantharidin | 9.62 | / |
| Temsirolimus | Rotenone | 9.60 | / |
| Temsirolimus | Azithromycin | 9.14 | / |
| Cantharidin | Enoxacin | 8.32 | / |

The "\*" symbol in the table highlights the drug combinations verified by experimental studies.

By searching databases such as PubChem and DrugBank(Knox et al., 2024), we identified the pathways associated with various drugs to understand the principles underlying drug combinations, thereby enhancing the interpretability of the prediction results. Here are examples of two types of drug combinations with different benefits: Combination with synergistic effects on the same pathway: For instance, the combination of Temsirolimus and Nitazoxanide (nitazoxanide). Both drugs inhibit the mTOR protein, thereby jointly blocking the PI3K-Akt signaling pathway, which corresponds to the pathways enriched by differential gene analysis of LAMA3/MET. Combination with synergistic effects on different pathways, for example, the combination of Fisetin and Azithromycin. Azithromycin reduces inflammation by inhibiting the NF-κB signaling pathway, corresponding to the NF-kappa B signaling pathway. Conversely, Fisetin upregulates HO-1 expression via the p38 MAPK pathway, inhibiting doxorubicin-induced senescence of pulmonary artery endothelial cells. It also inhibits the proliferation of pulmonary artery smooth muscle cells through the Nrf2/HO-1 signaling pathway, thereby preventing pulmonary artery remodeling. These correspond to the MAPK and Nrf2/HO-1 signaling pathways.

The specific relationships between the individual drugs and enriched pathways are detailed in Table 5.

**Table 5.**
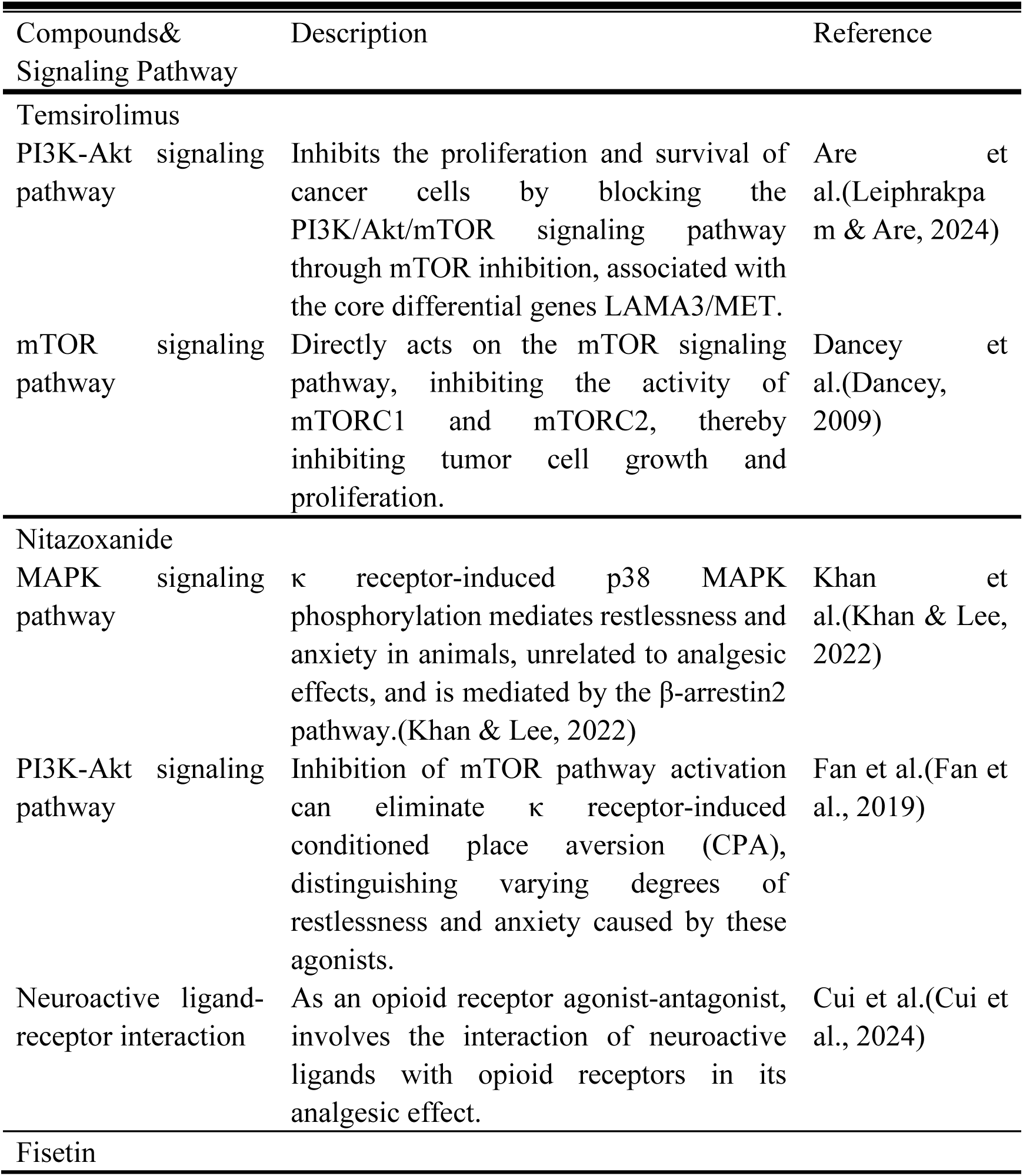
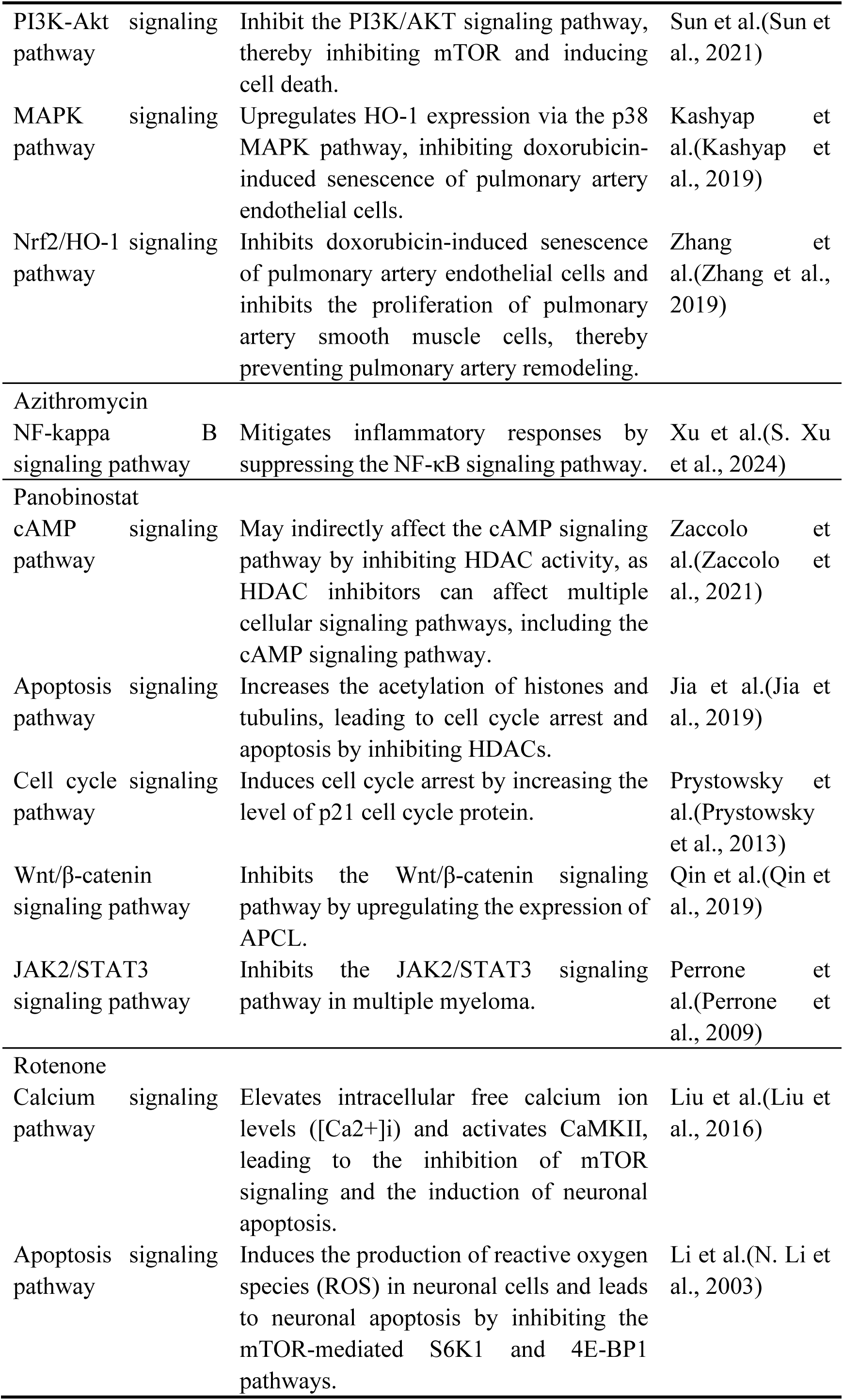
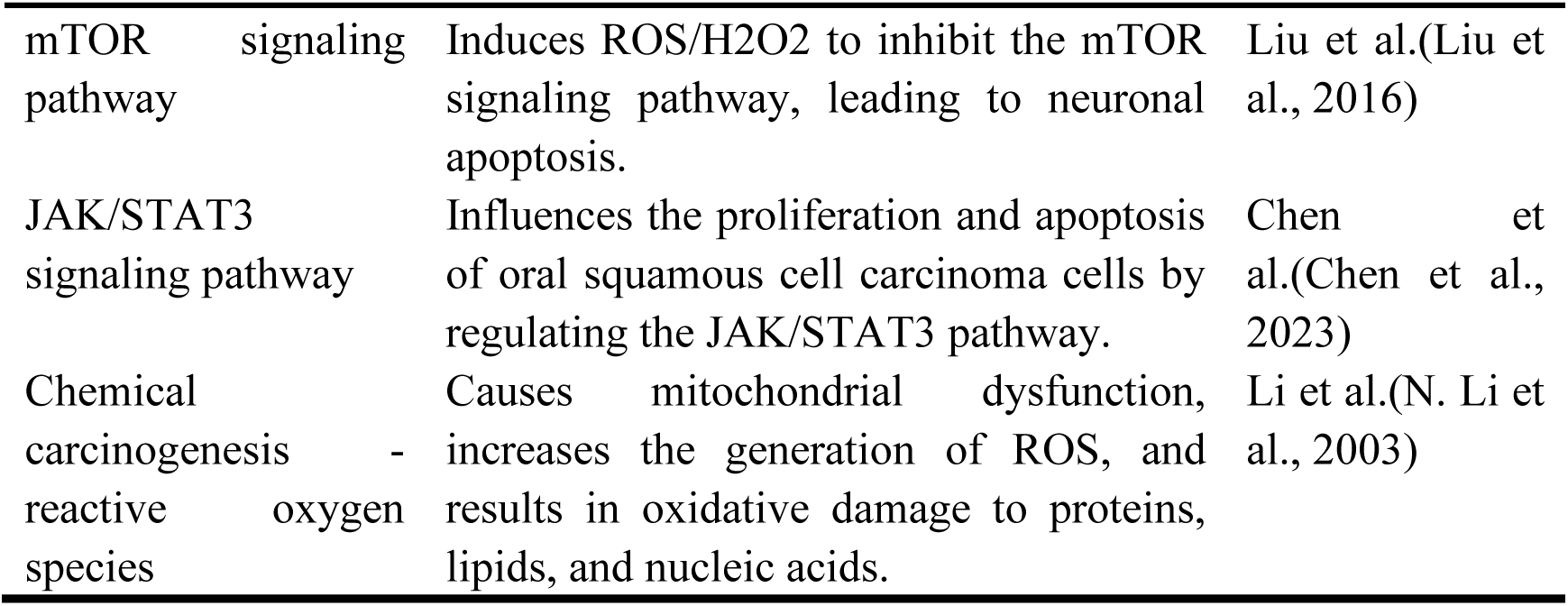
Senolytics and their associated signaling pathways.

### Verification

The inherent lack of interpretability in deep learning consistently poses challenges for model mechanism inference. In this study, we employed molecular docking simulations to validate the mechanisms of high-synergy drug combinations, enhancing the interpretability of synergistic effects in drug combinations and further elucidating the advantages of dual-drug systems. To validate the efficacy of our model, we conducted a literature review for the top ten predicted drug combinations. Based on the model’s scoring system, we identified a novel senolytic drug combination, Temsirolimus + Nitazoxanide, with the highest score. We performed molecular docking simulations to validate the interaction further and confirm its synergistic potential.

### Molecular docking verification

We simulated the docking of the highest-scoring drug combination predicted by the model: Temsirolimus+Nitazoxanide on the same target. The docking results indicated that the binding sites of these two compounds on the mTOR target were highly similar. In the context of the dynamic structure of the three-dimensional conformation of the protein, this drug combination demonstrates a broader binding range for FKBP12 in mTORC1 compared to Temsirolimus alone. Temsirolimus is a prodrug of rapamycin that is rapidly converted to its active form by cytochrome CYP 4503A4/5 in the bone marrow. It exhibits superior chemical stability and solubility compared with rapamycin. Its active component, rapamycin, is lipophilic and can permeate the cell membrane to bind to the intracellular receptor FKBP-12 (FK506-binding protein 12 kDa) (Oh et al., 2024). The resulting complex then binds to the FRB domain of the TOR protein, inhibiting its function.

We used AutoDock(Eberhardt et al., 2021) to complete the simulated docking of nitazoxanide with this protein, revealing a high consistency in the target action region for this drug combination. Upon adjusting the three-dimensional docking view of both compounds to the same angle as in Figure 5b), it becomes evident that the binding regions of the individual drugs within this combination highly overlap with that of mTOR. From the perspective of three-dimensional dynamic protein complementarity, this effectively demonstrates the expansion of the target hit range following drug combination therapy.

**Figure 4.**
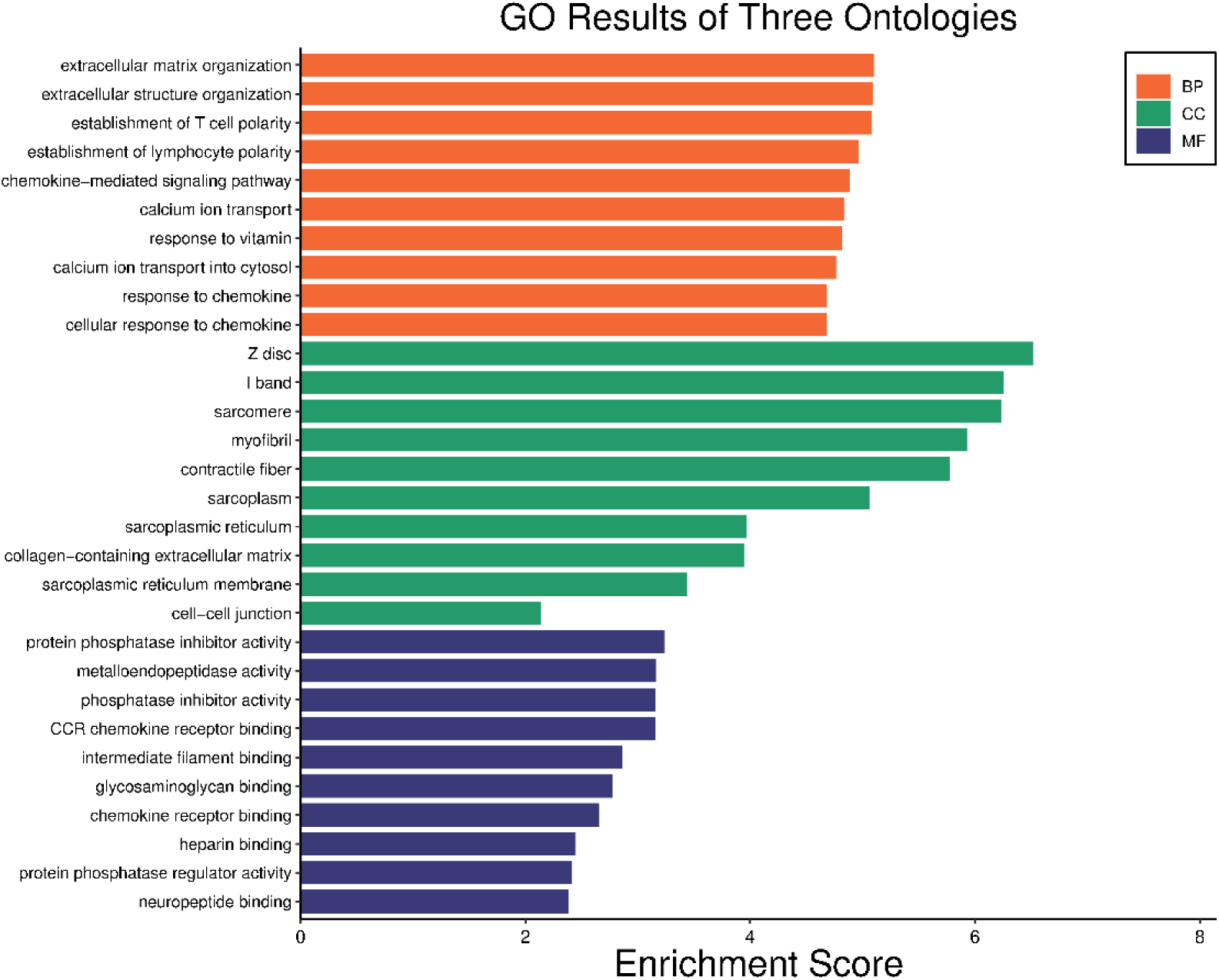
Senescence-related Differential Gene Enrichment Analysis. BP: The enrichment analysis of biological processes of aging-related core DEGs demonstrated their predominant involvement in extracellular matrix organization and extracellular structure organization. CC: The enrichment analysis of cellular components of aging- related core DEGs revealed a concentration of these genes in the Z disc, I band, and sarcomere, confirming their origin from bone tissue cells. MF: The enrichment analysis of molecular functions emphasized protein phosphatase inhibitor activity, metalloendopeptidase activity, phosphatase inhibitor activity, and CCR chemokine receptor binding, all closely associated with the aging process.

**Figure 5.**
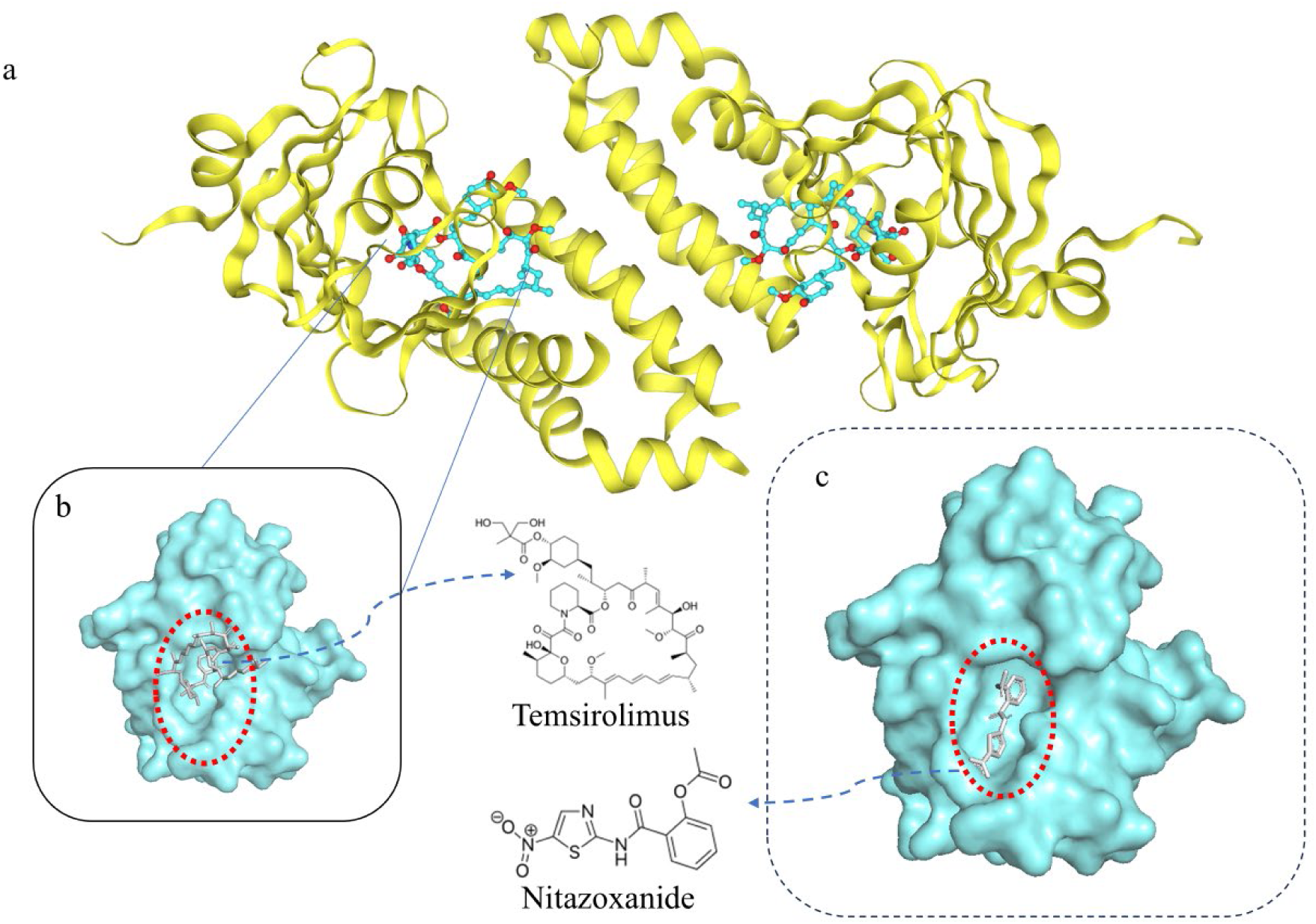
Molecular docking simulation verification of Temsirolimus and Nitazoxanide with the mTOR protein. 5a) shows the mTOR protein’s original docking site with the Temsirolimus’s active component, 5b) shows an enlarged view of the binding site between the active component of Temsirolimus and the mTOR protein, and 5c) displays the binding site after docking nitazoxanide to the mTOR protein.

### Literature verification

A comprehensive literature review revealed that certain combinations listed in Table 4 have been experimentally validated. For the combination of Cantharidin and Fisetin, which has a predicted synergistic score of 10.35, Frezzato and colleagues(Frezzato et al., 2020) substantiated its joint action in inhibiting the binding of HSF1 to the HSP70 promoter, resulting in the downregulation of HSP70 expression.

Similarly, the combined treatment of Canthardin and ABT-737 with a predicted synergistic score of 9.70 was explored by Ren and colleagues(Ren et al., 2016), who empirically demonstrated its enhanced inhibitory effect on cell proliferation, targeting Bcl-2 family proteins relevant to aging. Na and colleagues(Na et al., 2014) further confirmed through in vitro experiments the synergistic induction of apoptosis in cervical cancer cells with the combined application of norcantharidin and ABT-737, where NCTD significantly augmented the effect of ABT-737. Empirical evidence from the literature also highlighted the structural analog of baicalein, norcantharidin, in combination with ABT-737, elucidating a mechanism involving the inhibition of Mcl-1 through transcriptional suppression by NCTD, ultimately enhancing ABT-737- induced apoptosis in liver cancer cells. Norcantharidin and Cantharidin share similarities in chemical structure. Both compounds contain multiple cyclic structures and functional groups. Specifically, they both contain seven-membered and five-membered rings, which are connected by a carbonyl group. In addition, both include two carboxyl groups in similar positions on the ring.

The specific action mechanisms involve NCTD promoting the mitochondrial translocation of Parkin, leading to changes in the mitochondrial membrane potential and an increase in mitochondrial reactive oxygen species (ROS), as well as inducing the accumulation of autophagic vacuoles and blocking autophagic flux, thereby regulating the expression of apoptosis-related proteins.

The combination of Finasteride and Azithromycin has been shown to have a predicted synergistic score of 9.64. Shao and colleagues(Shao et al., 2024) also confirmed that the combination of fisetin, a member of the flavonoid family, and azithromycin exhibits a more potent anti-inflammatory effect. The specific mechanism involves inhibiting phosphorylation in the JAK/STAT and MAPK pathways and the nuclear translocation of NF-κB p65, alleviating tubal factor infertility (TFI). Notably, fisetin is homologous to quercetin, with a molecular difference of a hydroxyl group at the C5 position.

## CONCLUSION

This study, which uses differential genes associated with aging as the core entry point, thoroughly investigates the intricate relationships between senolytic drugs and apoptosis-related pathways. By integrating multi-omics analysis with advanced deep learning models, we have successfully applied a multi-drug combination prediction model that specifically evaluates the efficacy of senescent cell clearance. Based on this robust foundation, we have constructed a high-confidence senolytic drug combination database for predictive purposes, led by the combination of temsirolimus and nitazoxanide. The predicted results have been partially validated through extensive experimental literature data and molecular docking simulation calculations. This work provides a novel perspective on the diversity of clinical drug use, offers a new possible solution to the problem of limited combinatorial therapies in the clinical application of senolytic drugs, and may provide valuable insights into personalized aging prevention strategies.

## SUPPLEMENTARY INFORMATION

Additional file 1 of drug combination synergy prediction score using machine learning.

| drugA_name | drug_B_name | pred_syn_score |
| --- | --- | --- |
| Temsirolimus | Nitazoxanide | 13.28 |
| Temsirolimus | Fisetin | 11.86 |
| Cantharidin | Fisetin | 10.35 |
| Fisetin | Enoxacin | 9.89 |
| Cantharidin | ABT-737 | 9.70 |
| Fisetin | Azithromycin | 9.64 |
| Panobinostat | Cantharidin | 9.62 |
| Temsirolimus | Rotenone | 9.60 |
| Temsirolimus | Azithromycin | 9.14 |
| Cantharidin | Enoxacin | 8.32 |
| Cantharidin | Adapalene | 7.93 |
| Cantharidin | Nitazoxanide | 7.88 |
| Fisetin | Adapalene | 7.65 |
| Temsirolimus | Roxithromycin | 7.57 |
| Dasatinib | Cantharidin | 7.43 |
| Temsirolimus | Curcumin | 7.37 |
| Temsirolimus | ELTROMBOPAG | 7.26 |
| Curcumin | Fisetin | 7.24 |
| Tanespimycin | Enoxacin | 7.11 |
| Rotenone | Enoxacin | 7.03 |
| Fisetin | Nitazoxanide | 6.97 |
| Panobinostat | Enoxacin | 6.78 |
| Temsirolimus | cyclosporin A | 6.68 |
| Temsirolimus | Adapalene | 6.42 |
| Temsirolimus | Geldanamycin | 6.42 |
| Cantharidin | Geldanamycin | 6.08 |
| Panobinostat | Fisetin | 6.01 |
| Cantharidin | Curcumin | 5.94 |
| PF-573228 | Fisetin | 5.90 |
| Cantharidin | Rotenone | 5.75 |
| PF-573228 | Cantharidin | 5.53 |
| Curcumin | Enoxacin | 5.43 |
| Azithromycin | Nitazoxanide | 5.37 |
| Rotenone | Adapalene | 5.35 |
| Azithromycin | Enoxacin | 5.25 |
| ABT-737 | Fisetin | 5.24 |
| Nitazoxanide | Enoxacin | 5.22 |
| Cantharidin | Tanespimycin | 5.12 |
| Fisetin | Roxithromycin | 4.65 |
| Azithromycin | ELTROMBOPAG | 4.25 |
| Cantharidin | Amiloride hydrochloride | 4.17 |
| Azithromycin | Adapalene | 4.17 |
| Fisetin | ELTROMBOPAG | 3.79 |
| Fisetin | Amiloride hydrochloride | 3.68 |
| PF-573228 | Enoxacin | 3.58 |
| Temsirolimus | Tanespimycin | 3.44 |
| Panobinostat | Curcumin | 3.30 |
| Rotenone | cyclosporin A | 3.28 |
| Rotenone | Nitazoxanide | 3.21 |
| Curcumin | Nitazoxanide | 3.17 |
| Rotenone | Azithromycin | 3.16 |
| Temsirolimus | Enoxacin | 3.11 |
| Cantharidin | Azithromycin | 3.10 |
| Panobinostat | Temsirolimus | 3.03 |
| Rotenone | ELTROMBOPAG | 2.99 |
| Temsirolimus | Digoxin | 2.96 |
| Curcumin | ELTROMBOPAG | 2.87 |
| ABT-737 | Adapalene | 2.84 |
| Cantharidin | ELTROMBOPAG | 2.82 |
| ABT-737 | Nitazoxanide | 2.81 |
| Curcumin | Roxithromycin | 2.75 |
| Cantharidin | Roxithromycin | 2.63 |
| Adapalene | Enoxacin | 2.56 |
| Fisetin | Rotenone | 2.56 |
| Curcumin | Azithromycin | 2.54 |
| Temsirolimus | Amiloride hydrochloride | 2.50 |
| Amiloride hydrochloride | Enoxacin | 2.44 |
| Adapalene | Roxithromycin | 2.41 |
| ABT-737 | Enoxacin | 2.34 |
| Enoxacin | ELTROMBOPAG | 2.10 |
| Adapalene | Nitazoxanide | 2.06 |
| PF-573228 | Temsirolimus | 2.02 |
| PF-573228 | Adapalene | 2.01 |
| Dasatinib | Enoxacin | 1.73 |
| Digoxin | Enoxacin | 1.72 |
| PF-573228 | Azithromycin | 1.61 |
| Panobinostat | Azithromycin | 1.47 |
| Roxithromycin | Enoxacin | 1.45 |
| Cantharidin | Digoxin | 1.42 |
| Nitazoxanide | ELTROMBOPAG | 1.39 |
| Dasatinib | Fisetin | 1.31 |
| Curcumin | cyclosporin A | 1.21 |
| ABT-737 | Curcumin | 1.17 |
| Panobinostat | Adapalene | 1.06 |
| Rotenone | Amiloride hydrochloride | 0.88 |
| Fisetin | cyclosporin A | 0.87 |
| PF-573228 | Curcumin | 0.84 |
| Roxithromycin | ELTROMBOPAG | 0.82 |
| PF-573228 | ELTROMBOPAG | 0.75 |
| Azithromycin | Amiloride hydrochloride | 0.59 |
| Dasatinib | ELTROMBOPAG | 0.58 |
| ABT-737 | Azithromycin | 0.38 |
| Enoxacin | cyclosporin A | 0.29 |
| Azithromycin | Roxithromycin | 0.21 |
| Curcumin | Digoxin | 0.20 |
| PF-573228 | ABT-737 | 0.02 |
| Dasatinib | Azithromycin | -0.04 |
| Adapalene | ELTROMBOPAG | -0.20 |
| Curcumin | Rotenone | -0.24 |
| Panobinostat | Nitazoxanide | -0.25 |
| ABT-737 | Temsirolimus | -0.25 |
| Panobinostat | ELTROMBOPAG | -0.25 |
| Nitazoxanide | cyclosporin A | -0.34 |
| Azithromycin | cyclosporin A | -0.41 |
| Tanespimycin | Adapalene | -0.43 |
| Rotenone | Roxithromycin | -0.44 |
| Nitazoxanide | Amiloride hydrochloride | -0.50 |
| Panobinostat | ABT-737 | -0.56 |
| Geldanamycin | Enoxacin | -0.58 |
| PF-573228 | Nitazoxanide | -0.60 |
| Adapalene | cyclosporin A | -0.68 |
| Dasatinib | Temsirolimus | -0.69 |
| Cantharidin | cyclosporin A | -0.78 |
| Fisetin | Tanespimycin | -0.82 |
| Curcumin | Amiloride hydrochloride | -0.84 |
| Curcumin | Adapalene | -0.88 |
| Nitazoxanide | Roxithromycin | -1.10 |
| Tanespimycin | Azithromycin | -1.15 |
| Amiloride hydrochloride | ELTROMBOPAG | -1.22 |
| ABT-737 | Amiloride hydrochloride | -1.39 |
| Cantharidin | Temsirolimus | -1.45 |
| Dasatinib | Curcumin | -1.63 |
| Fisetin | Digoxin | -1.65 |
| ABT-737 | cyclosporin A | -1.78 |
| PF-573228 | cyclosporin A | -1.85 |
| Curcumin | Geldanamycin | -1.87 |
| ABT-737 | Roxithromycin | -1.90 |
| Dasatinib | PF-573228 | -1.92 |
| Panobinostat | cyclosporin A | -1.94 |
| Tanespimycin | Rotenone | -1.95 |
| Roxithromycin | cyclosporin A | -1.97 |
| Digoxin | Nitazoxanide | -2.01 |
| Digoxin | ELTROMBOPAG | -2.02 |
| Tanespimycin | Nitazoxanide | -2.09 |
| ABT-737 | ELTROMBOPAG | -2.10 |
| Fisetin | Geldanamycin | -2.35 |
| PF-573228 | Roxithromycin | -2.42 |
| Panobinostat | Roxithromycin | -2.56 |
| PF-573228 | Amiloride hydrochloride | -2.86 |
| PF-573228 | Rotenone | -2.98 |
| Rotenone | Digoxin | -3.32 |
| Geldanamycin | Azithromycin | -3.33 |
| Digoxin | Roxithromycin | -3.38 |
| cyclosporin A | ELTROMBOPAG | -3.42 |
| Panobinostat | Amiloride hydrochloride | -3.46 |
| Amiloride hydrochloride | cyclosporin A | -3.49 |
| Digoxin | Adapalene | -3.49 |
| Digoxin | Azithromycin | -3.52 |
| Digoxin | cyclosporin A | -3.60 |
| Amiloride hydrochloride | Roxithromycin | -3.70 |
| Dasatinib | Nitazoxanide | -3.76 |
| ABT-737 | Rotenone | -3.84 |
| Tanespimycin | Roxithromycin | -3.90 |
| Curcumin | Tanespimycin | -4.19 |
| Geldanamycin | Adapalene | -4.21 |
| Dasatinib | Roxithromycin | -4.25 |
| PF-573228 | Panobinostat | -4.35 |
| Tanespimycin | ELTROMBOPAG | -4.41 |
| Panobinostat | Rotenone | -4.54 |
| Tanespimycin | Geldanamycin | -4.67 |
| Adapalene | Amiloride hydrochloride | -4.67 |
| Tanespimycin | cyclosporin A | -5.29 |
| Dasatinib | Amiloride hydrochloride | -5.43 |
| PF-573228 | Geldanamycin | -5.58 |
| ABT-737 | Digoxin | -5.76 |
| ABT-737 | Geldanamycin | -5.81 |
| Geldanamycin | ELTROMBOPAG | -5.83 |
| Tanespimycin | Amiloride hydrochloride | -6.15 |
| Dasatinib | Adapalene | -6.33 |
| Geldanamycin | Roxithromycin | -6.38 |
| Dasatinib | ABT-737 | -6.64 |
| Dasatinib | cyclosporin A | -6.87 |
| Geldanamycin | Nitazoxanide | -6.88 |
| PF-573228 | Digoxin | -6.92 |
| Dasatinib | Panobinostat | -7.03 |
| Panobinostat | Digoxin | -7.26 |
| Digoxin | Amiloride hydrochloride | -7.38 |
| Dasatinib | Geldanamycin | -8.06 |
| Geldanamycin | Digoxin | -8.20 |
| Geldanamycin | cyclosporin A | -8.48 |
| Dasatinib | Rotenone | -8.60 |
| Geldanamycin | Amiloride hydrochloride | -8.73 |
| Panobinostat | Geldanamycin | -9.52 |
| ABT-737 | Tanespimycin | -9.54 |
| PF-573228 | Tanespimycin | -9.82 |
| Panobinostat | Tanespimycin | -10.00 |
| Geldanamycin | Rotenone | -10.12 |
| Tanespimycin | Digoxin | -10.27 |
| Dasatinib | Tanespimycin | -11.78 |
| Dasatinib | Digoxin | -12.37 |

